# PINK1 is a target of T cell responses in Parkinson’s disease

**DOI:** 10.1101/2024.02.09.579465

**Authors:** Gregory P. Williams, Tanner Michaelis, João Rodrigues Lima-Junior, April Frazier, Ngan K. Tran, Elizabeth J. Phillips, Simon A. Mallal, Irene Litvan, Jennifer G. Goldman, Roy N. Alcalay, John Sidney, David Sulzer, Alessandro Sette, Cecilia S. Lindestam Arlehamn

## Abstract

Parkinson’s disease (PD) is associated with autoimmune T cells that recognize the protein alpha-synuclein in a subset of individuals. Multiple neuroantigens are targets of autoinflammatory T cells in classical central nervous system autoimmune diseases such as multiple sclerosis (MS). Here, we explored whether additional autoantigenic targets of T cells in PD. We generated 15-mer peptide pools spanning several PD-related proteins implicated in PD pathology, including GBA, SOD1, PINK1, parkin, OGDH, and LRRK2. Cytokine production (IFNγ, IL-5, IL-10) against these proteins was measured using a fluorospot assay and PBMCs from patients with PD and age-matched healthy controls. This approach identified unique epitopes and their HLA restriction from the mitochondrial-associated protein PINK1, a regulator of mitochondrial stability, as an autoantigen targeted by T cells. The T cell reactivity was predominantly found in male patients with PD, which may contribute to the heterogeneity of PD. Identifying and characterizing PINK1 and other autoinflammatory targets may lead to antigen-specific diagnostics, progression markers, and/or novel therapeutic strategies for PD.

## Introduction

Parkinson’s disease (PD) pathobiology is characterized by the formation of aggregated alpha-synuclein (α-syn) and subsequent neurodegeneration (1). This prominent role of α-syn in PD development is supported by reports that the gain of function genetic variation in the *SNCA* gene can be causal in rare inherited forms of PD (2) or increase one’s risk of developing idiopathic PD (3). However, several other genes and their proteins are associated with the pathology and development of PD. For example, perturbations in *PINK1* (4), *PRKN* (5), *GBA* (6), and *LRRK2* (*7*) have each been discovered to be major genetic risk factors for the development of PD. Multiple proposed mechanisms surrounding their pathogenicity role include dysfunctional autophagy (8) and/or mitophagy (9).

Recently, these PD-related proteins have also been linked to the activation of the immune system (10–12). Initial studies provided evidence that infiltrating T cells could be found in postmortem PD brain parenchyma (13, 14), but the role of these cells was unclear. Our group has shown that some PD patients harbor T cells recognizing α-synuclein (α-syn) (15–17). Furthermore, another group recently showed that these synuclein-specific T cells are associated with neurodegenerative signaling in PD and Lewy Body dementia, a related synucleinopathy (18). The presence of the α-syn-specific T cells correlates with the PD duration (16, 17). However, not all patients with PD have α-syn-specific T cells, even at early time points following diagnosis, leading us to hypothesize that additional autoantigens may exist. For example, in other diseases such as multiple sclerosis, several myelin-derived antigens are targeted by auto-inflammatory T cells (19). With this rationale, we screened a cohort of individuals with PD and age-matched healthy controls using peptide pools targeting several PD-associated proteins (either implicated genetically or in preclinical disease models) and measured the resulting T cell responses. Here, we report that PD patients have more frequent responses with a higher magnitude of cytokine production towards the mitochondrial-associated protein PINK1. This difference was driven by male PD patients, providing a sex-specific difference in antigen recognition. We further report the identification of specific PINK1 epitopes mediating the auto-antigenic T cell response. These findings indicate additional antigenic targets in PD and emphasize the promise for potential immune-based biomarkers and therapies in its treatment.

## Results

### Screening PD-related proteins for autoantigenic T cell responses

Peripheral blood mononuclear cells (PBMCs) were processed from whole-blood donations provided by individuals with PD (n=39) and healthy controls (HC; n=39). Participants of the study were recruited from three different sites across the US: New York (Columbia University Irving Medical Center), Illinois (Shirley Ryan AbilityLab/Northwestern University), and California (University of California San Diego and La Jolla Institute for Immunology). Detailed cohort demographics, including the clinical characteristics of the PD cohort, can be found in **Supplemental Table 1**. We tested six PD-related proteins as potential targets of T cell recognition in individuals with PD. These proteins were selected based on their genetic link to PD, their presence in Lewy bodies, and/or being implicated in preclinical models of PD: (PINK1 (4, 11, 12, 20, 21), parkin (5, 11, 12), OGDH (11, 22), GBA (6, 8, 23), SOD1 (24, 25), and LRRK2 (7, 21) (**Fig. 1a**).

**Figure 1:**
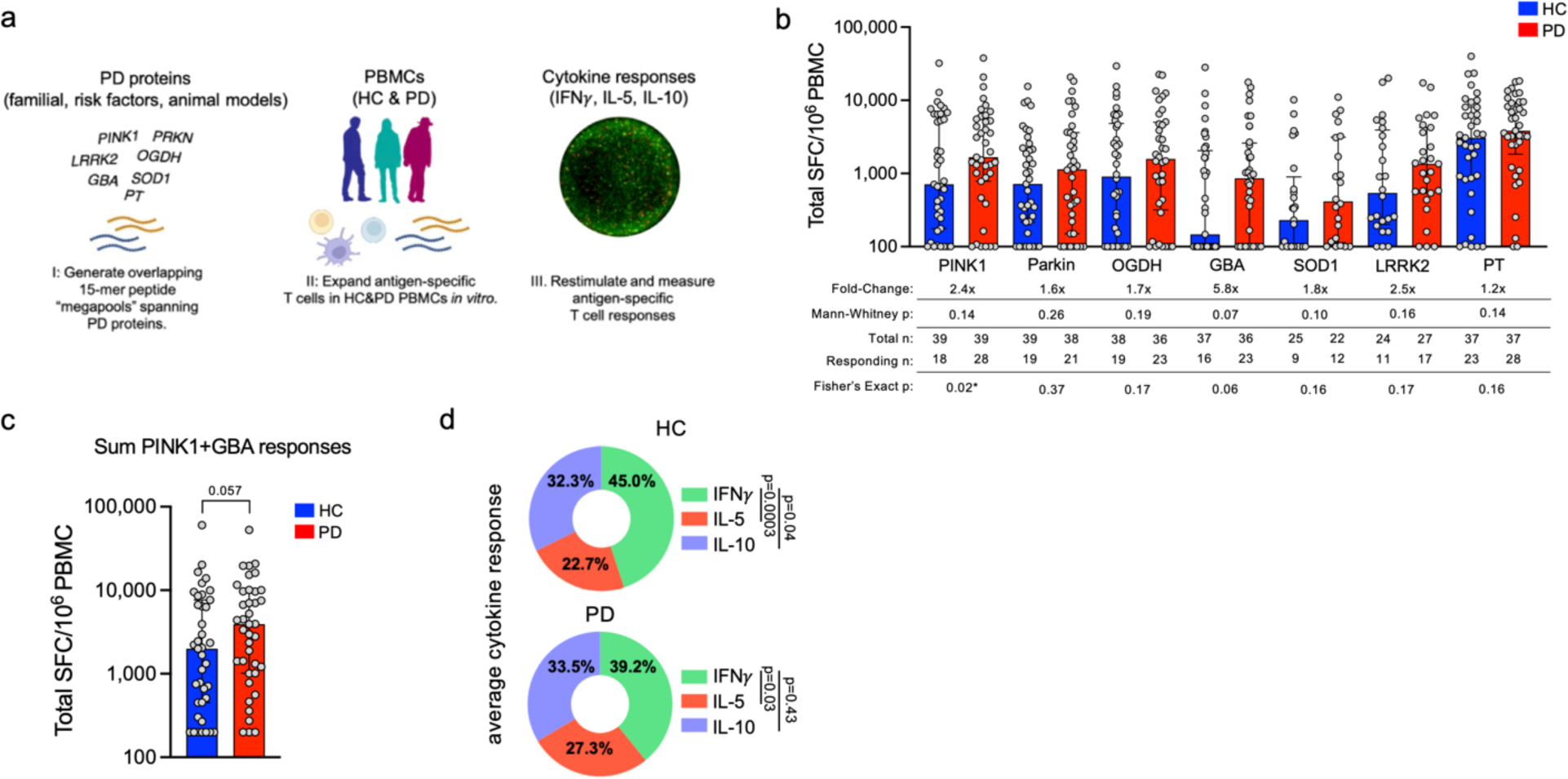
Screening PD-related proteins for autoantigenic T cell responses. a) Experimental design for the screening of PD related proteins. I) 15-mer peptides spanning PD-related proteins; PINK1 (117 peptides), PARKIN (94 peptides), OGDH (203 peptides), GBA (106 peptides), SOD1 (34 peptides), LRRK2 (80 predicted peptides), and PT as a control (132 peptides). II) Peptide pools were incubated at a concentration of 5 ug/mL with PBMCs from PD participants and age-matched HC for 14 days. III) Restimulation of cultured PBMCs with the initial antigen pools and subsequent determination of antigen-specific cytokine production using Fluorospot. DMSO and PHA stimuli were used as negative and positive controls, respectively, for each participant/pool combination. b) Magnitude of total cytokine response (sum of IFN*γ*, IL-5, and IL-10) to neuroantigens and control PT between HC (blue bars) and PD (red bars), each circle representing an individual participant. Median ± interquartile range displayed. Fold-change is in comparison to HC response. One-tailed Mann-Whitney tests were performed between HC and PD antigen-cytokine values. One-tailed Fisher tests performed using the geometric mean of the HC group for each individual antigen as a cutoff for the test. PINK1(PD n=39, HC n=39), PARKIN (PD n=37, HC n=39), OGDH (PD n=36, HC n=38), GBA (PD n=37, HC n=36), SOD1 (PD n=24, HC n=25), LRRK2 (PD n=26, HC n=24), and PT as a control (PD n=37, HC n=37) c) Summed cytokine responses towards PINK1 and GBA antigen pools between HC and PD participants. Median ± interquartile range displayed. One-tailed Mann-Whitney test. d) Average % cytokine of total response (i.e., IFN6FE;/sum of IFN6FE;/IL-5/IL-10) between HC and PD across all neuroantigens tested (PINK1, PARKIN, OGDH, GBA, SOD1, and LRRK2). One-way ANOVA with Dunnett’s test.

To determine whether T cells recognize these PD-related proteins, we assayed pools of 15 amino acid peptides overlapping by ten residues and spanning the full sequence of each protein; PINK1 (117 peptides), parkin (94 peptides), OGDH (203 peptides), GBA (106 peptides), SOD1 (34 peptides), and peptides predicted to bind HLA class II alleles for LRRK2 (80 peptides). Individual peptide sequences and more detailed pool information can be found in **Supplemental Table 2**. As a control, we also included a previously described peptide pool directed towards *Bordetella pertussis* vaccine antigens (26) (PT, **Supplemental Table 2**), which individuals are exposed to through Tdap vaccination. We hypothesized that there would be a higher magnitude of T cell-specific responses against these antigens in individuals with PD compared to age-matched HC. PBMCs from the PD and HC cohorts were stimulated *in vitro* with the different peptide pools for 14 days. At the end of the restimulation period, expanded cultures were assayed by tri-color Fluorospot, measuring IFNγ, IL-5, and IL-10 (**Fig. 1a**), which were selected as representative of Th1, Th2 and Treg responses, respectively.

### Higher PINK1-specific T cell reactivity in PD patients compared to controls

The T cell reactivity to the neuroantigen pools is shown in **Fig. 1b**. A significant increase in the frequency of PINK1 reactivity was observed among PD patients compared to HC along with a trend for an increased magnitude of the PINK1 response (2.4-fold increase, Fisher’s exact one-tailed test, p=0.02; one-tailed Mann-Whitney, p=0.14).

The response magnitude of the PD patients was also higher than those observed in the HC cohort for other neuroantigens. However, the increases did not reach statistical significance; 5.8-fold in the case of GBA (M-W p=0.07, Fisher’s p=0.06), 2.5-fold in the case of LRRK2 (M-W p=0.16, Fisher’s p=0.17), 1.8-fold in the case of SOD1 (M-W p=0.10, Fisher’s p=0.16), 1.7-fold in the case of OGDH (M-W p=0.19, Fisher’s p=0.17), and 1.6-fold in the case of Parkin (M-W p=0.26, Fisher’s p=0.37). Responses to the control PT peptide pool were 1.2-fold increased.

When the overall responses to the PINK1 and GBA antigens were considered in aggregate, we observed a 2.0-fold increase in the magnitude of response in PD patients compared to HC (**Fig. 1c,** p=0.057 one-tailed Mann Whitney). The individual cytokine profiles were highly polyfunctional. When the response to all neuroantigens was considered in aggregate, IFNγ accounted for 39.2% of the total, IL-5 for 27.3%, and IL-10 for 33.5% in PD patients with a similar profile in HC (**Fig 1d**). IFNγ responses were significantly more prevalent than IL-5 for both HC and PD (p=0.003 and 0.03 respectively, one-way ANOVA with Dunnett’s test), and only in HC were significantly higher than IL-10 (p=0.04 HC, p=0.43 PD; one-way ANOVA with Dunnett’s test). Detailed results for the individual cytokines compared between HC and PD participants are in **Supplemental Fig. 1.** In conclusion, PINK1-specific T cell responses and potentially additional neuroantigen-specific responses are higher in PD patients than in HC.

### T cell reactivity in PD is not associated with early time points or other clinical characteristics

We examined the correlation between neuroantigen-specific T cell reactivity and disease status, including age, time from diagnosis, cognitive function (the Montreal Cognitive Assessment (MoCA) (27), motor examination (Part III from the Unified Parkinson’s Disease Rating Scale (UPDRS) (28), and medication (levodopa equivalent dose; LED (29)) scores. We only found a positive correlation between age and T cell reactivity to LRRK2, a negative correlation between time since diagnosis (years) and T cell reactivity to GBA and SOD1, and between LED and T cell reactivity to SOD1 (**Supplemental figure 2**). No correlations between these parameters and T cell reactivity to PINK1 were found. We previously found that the α-syn-specific T cell reactivity was higher closer to PD diagnosis and then waned (16), similar to the observations found here for GBA and SOD1. Thus, neuroantigen-specific T cell reactivity is complex, and the responses to the candidate antigens are differentially affected by age and time from diagnosis.

### Neuroantigen-specific T cell responses as a function of biological sex

It is well established that the incidence of PD is higher in males versus females (30). We observed that the increased PINK1 response in PD appeared predominantly driven by differences in males (**Fig. 2a,b)** with a 5.1-fold increase of PD vs. HC in males (Mann-Whitney p=0.09, Fisher’s exact p=0.02) compared to a 1.2-fold difference in female PD vs HC (Mann-Whitney p=0.23, Fisher’s exact p=0.63). Similarly, LRRK2 responses had a trend for higher magnitudes in PD vs. HC males (5.8-fold, Mann-Whitney p=0.07, Fisher’s exact p=0.09) versus females (0.8-fold difference in PD vs. HC; Mann Whitney p=0.38, Fisher’s exact p=0.61). A similar trend was noted for parkin (2.1-fold for PD vs. HC males, Mann-Whitney p=0.20, Fisher’s exact p=0.24) versus females having a 0.8-fold difference in PD vs. HC (Mann-Whitney p=0.15, Fisher’s exact p=0.44). The SOD1-specific response magnitude trended higher in PD females compared to HC females (**Fig. 2b**, 4.1-fold increase, Mann Whitney p=0.07, Fisher’s exact p=0.13), as did GBA reactivity (4.6-fold increase, Mann Whitney p=0.32, Fisher’s exact p=0.17).

**Figure 2:**
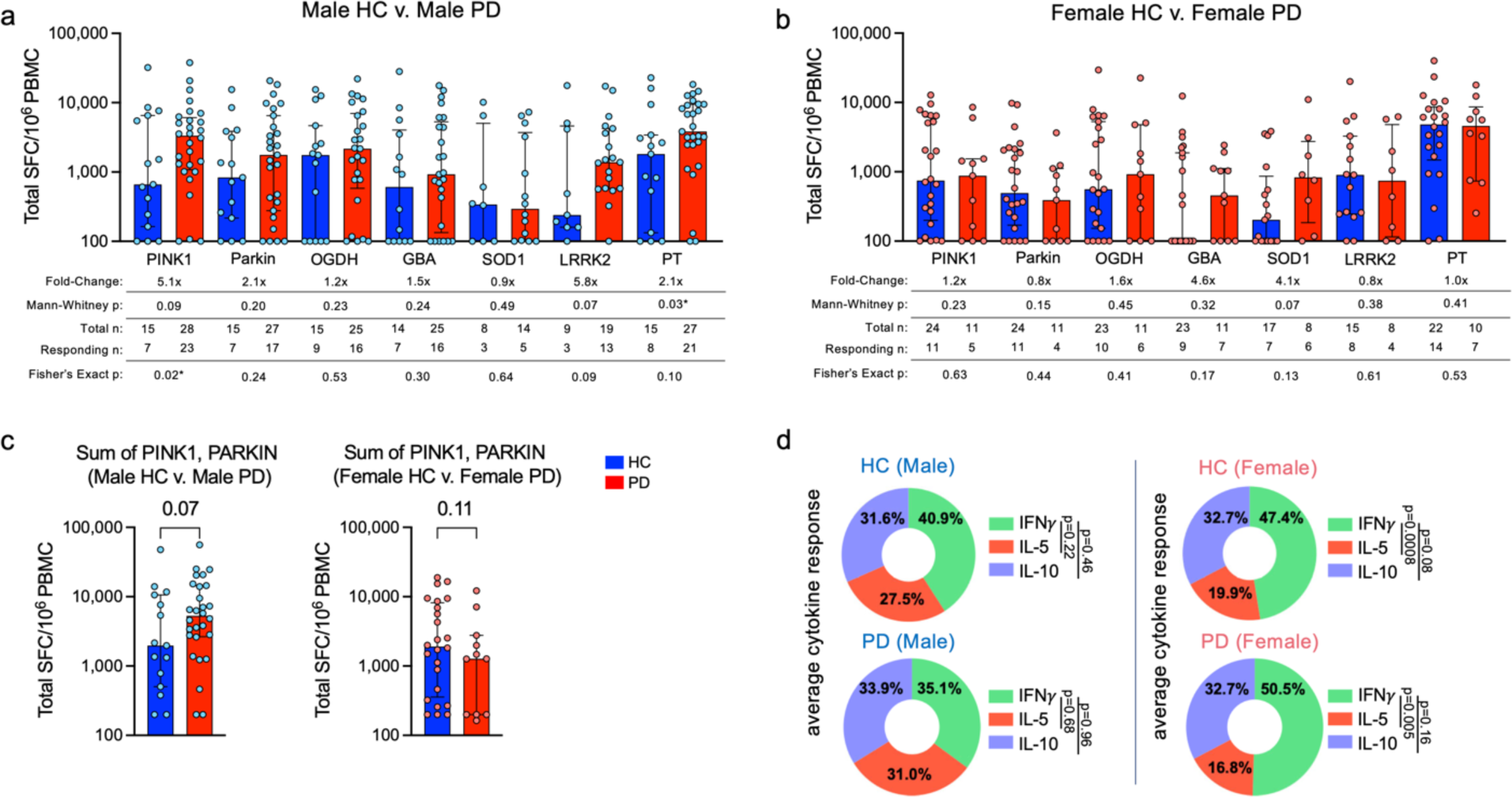
Neuroantigen-specific T cell responses as a function of biological sex. Magnitude of total cytokine response (sum of IFN6FE;, IL-5, and IL-10) to neuroantigens and control PT between a) male HC and PD; b) female HC and PD. HC (blue bars) and PD (red bars), each circle representing an individual participant. Median ± interquartile range displayed. Fold-change is in comparison to HC response. One-tailed Mann-Whitney tests were performed between HC and PD antigen-cytokine values. One-tailed Fisher tests were performed using the geometric mean of the HC group for each individual antigen as a cutoff for the test. c) Summed cytokine responses towards PINK and PARKIN antigens between male HC/PD (left panel) and female HC/PD (right panel). Median ± interquartile range displayed, One-tailed Mann-Whitney test. d) Average % cytokine of total response (i.e. IFN6FE;/sum of IFN6FE;/IL-5/IL-10) between HC and PD across all neuroantigens tested (PINK1, PARKIN, OGDH, GBA, SOD1, and LRRK2). One-way ANOVA with Dunnett’s test.

PINK1 and parkin are thought to work in tandem to handle mitochondrial turnover (31, 32). We compared the sum of reactivity to PINK1 and parkin in individual male PD vs. HC participants to reactivity in females, and we found a similar trend for an increased response towards these mitochondrial-associated proteins in males with PD (**Fig. 2c**, Mann-Whitney p=0.07), but not in females with PD (**Fig. 2c**, Mann-Whitney p=0.11). When the total responses of male vs. female participants were broken down into their individual cytokine constituents, we observed comparable responses as reported above for the entire cohort (**Fig. 1d**). IFNγ was still the most prominent cytokine produced against all antigens, with males (40.9% male HC, 35.1% male PD) not significantly different from females (47.4% female HC, 50.5% in female PD; **Fig. 2d**). IL-5 accounted for 27.5% and 31.0% of the cytokine response in male HC and PD, and in female HC and PD, was 19.9% and 16.8% respectively. The IL-10 response in male HC and PD response was 31.6% and 33.9%, respectively, compared to female HC and PD (both 32.7%, **Fig. 2d**). Intriguingly, it appears that the IFNγ bias towards neuroantigens is much more pronounced in both female HC and PD compared to males. In female HC and PD, but not male, IFNγ response was significantly higher than IL-5 (p=0.0008 and 0.005 respectively, one-way ANOVA with Dunnett’s test) and trended higher in comparison to IL-10 (p=0.08 female HC, p=0.16 female PD; one-way ANOVA with Dunnett’s test). More detailed individual antigen/cytokine differences between male and female HC and PD can be found in **Supplemental Fig. 3**. Taken together, these results suggest that reactivity to different PD autoantigens have a sex bias in terms of specific antigen reactivity and the types of cytokines produced in response.

### Phenotypic characterization of PINK1 responsive T cells

We then characterized in more detail the phenotype of the expanding/cytokine-producing cells in cultures stimulated with the PINK1 peptide pool. After *in vitro* expansion, we analyzed PBMC cultures from a subset of PD participants (n=6, 5 males, 1 female) stimulated with the PINK1 peptide pool by flow cytometry (**Fig. 3a**). Gating on live, single, CD3^+^ cells (**Fig. 3b**), the predominant cell type in the PINK1 expanded cultures was CD4^+^ T cells (64% ± 18% of live cells, **Fig. 3c**), which were present in significantly higher frequencies than CD8^+^ T cells (16% ± 8%, p<0.0001) and non-CD3 cells (15% ± 9%, p<0.0001). This data demonstrates that the predominant cell type recognizing the PINK1 epitopes is CD4+ T cells, consistent with what has been observed for other PD neuroantigens (16, 17, 33).

**Figure 3:**
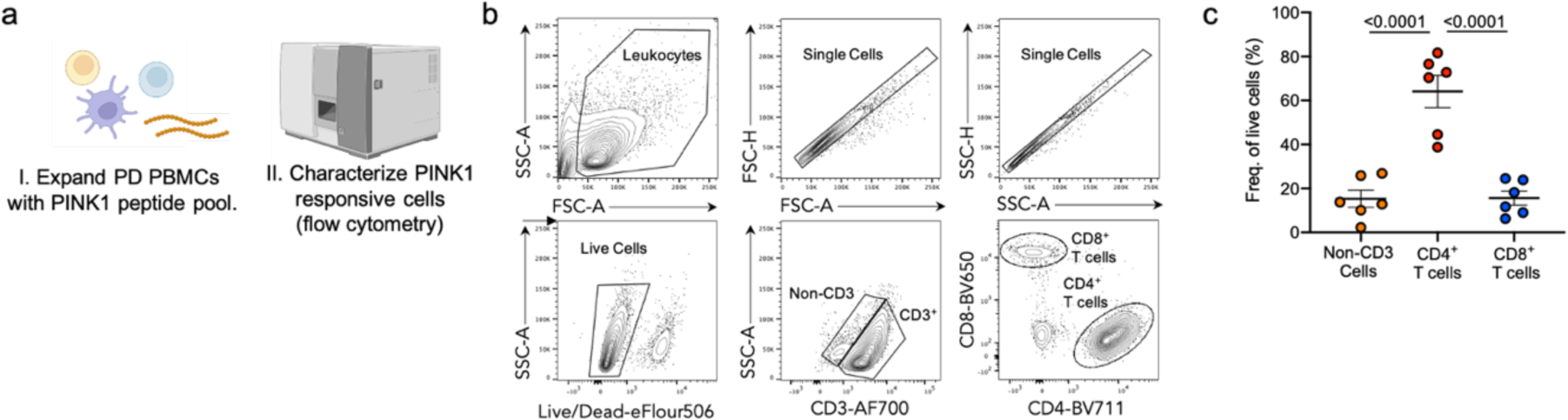
Phenotypic characterization of PINK1 responsive T cells. a) Diagram describing experimental design to characterize PINK1 expanded PBMC cultures from individuals with PD. b) Representative gating strategy depicting the identification and quantification of live, singlet, non-CD3^+^ or CD3^+^, CD4^+^/CD8^+^ T cells. c) Frequency of CD4 (red circles), CD8 (blue circles), and non-CD3 (orange circles)) from PINK1 stimulated PD PBMCs. One-way ANOVA with Tukey’s multiple comparisons, mean ± SEM displayed.

### Identification of individual PINK1 epitopes eliciting T cell responses in PD

To identify individual PINK1 epitopes, we stimulated a subset (n=18; 15 male and 3 female) of previously identified PD PINK1 responders with the pool of PINK1 overlapping peptides. The resulting cultures were re-stimulated with 10 separate PINK1 “mesopools” (smaller pools of ∼12 individual peptides) that spanned the PINK1 protein. The top 3 highest mesopool responses for each participant were then deconvoluted to identify individual PINK1 epitopes (**Fig. 4a**). We identified 34 individual peptides that elicit T cell responses in PD patients (**Fig. 4b, Supplemental Table 2**). The average number of PINK1 epitopes recognized by each PD patient was 5.2 (median of 5.5, range 1-14, **Fig. 4c)**. The dominant epitope, aa216 LAIKMMWNISAGSSS, was recognized by 55.5% of the PD patients (**Fig. 4d**). The next most recognized epitope, aa220 APAFPLAIKMMWNIS, was recognized in 27.7% of PD patients. A total of seven epitopes were recognized in 3 or more participants. Of the three female PD patients included in these experiments, 2/3 had T cell responses against the dominant aa216 LAIKMMWNISAGSSS epitope, while the remaining participant’s dominant epitope was aa511 LWGEHILALKNLKLD (also observed in a male participant).

**Figure 4:**
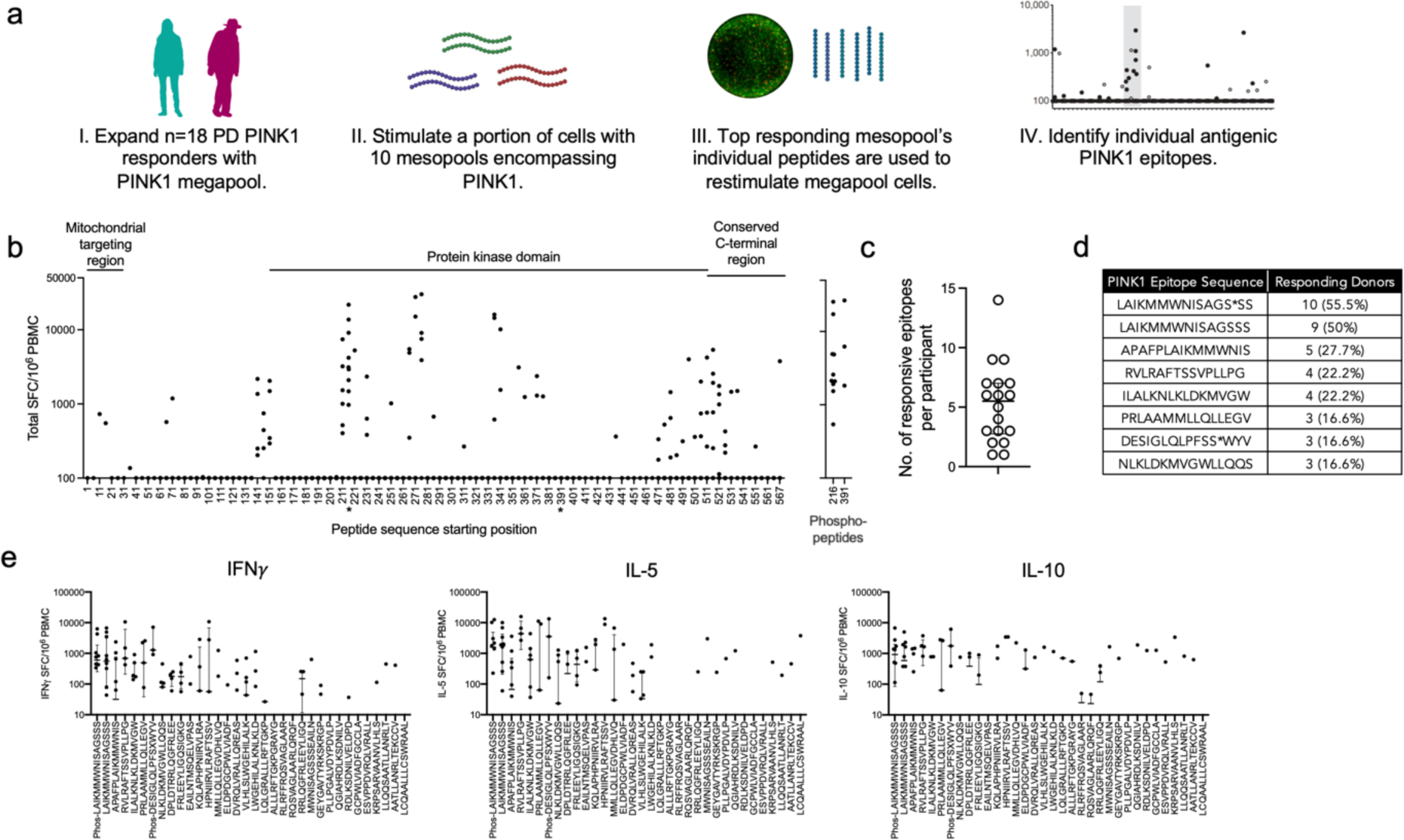
Identification of PINK1 epitopes eliciting T cell responses in PD. a) Experimental design utilized to identify PINK1 epitopes. PINK1 megapool was used to expand previously identified PD PINK1 responders. A portion of megapool expanded cells for each participant were then re-stimulated with 10 PINK1 mesopools (smaller pools containing, on average, 12 individual PINK1 epitopes). The individual epitopes from the top 3 responding mesopools for each participant were then used to restimulate the remaining megapool expanded cells, allowing for the identification of individual antigenic PINK1 epitopes. b) Individual PINK1 epitope responses (total cytokine, sum of IFN6FE;, IL-5, IL-10) displayed in relation to the major regions of the PINK1 protein (left to right across amino acid 1—581; * indicates peptides that were also included as phosphorylated versions). The right graph displays phosphorylated peptides. Each dot is a participant/peptide combination. c) Number of individual epitopes recognized by each of the 18 individual PD participants tested. d) Table displaying the identity and frequency of responses towards most commonly recognized PINK1 epitopes. * Indicates a phospho-serine at that position. e) Individual cytokine responses (IFN6FE;, IL-5, and IL-10) towards the 34 PINK1 epitopes displayed in order of frequency of recognition. Median ± interquartile range is shown.

Individual IFNγ, IL-5, and IL-10 responses towards the 34 identified PINK1 epitopes showed a similar pattern of responses as the original PINK1 megapool, with all three cytokines represented (**Fig. 4e).** Of note, the most commonly recognized epitope (i.e., a.a.216LAIKMMWNISAGSSS) typically elicited responses with all three cytokines with some participants producing all three cytokines against the epitope, while others only one or two. As expected, some more unique, single-participant responsive epitopes only resulted in one specific cytokine (IL-5 in the case of aa567 LCQAALLLCSWRAAL).

### Determination of potential HLA restriction of PINK1 epitopes

The results above indicate that CD4 T cells are the predominant T cell subset expanded following PINK1 peptide pool stimulation. To infer potential HLA restrictions, we examined each epitope recognized in two or more participants, following an approach outlined previously (34) and the NetMHCIIpan EL 4.1 tool hosted in the IEDB analysis resource. Inferred restrictions indicated by the corresponding beta chain are summarized in **Table 1**. For each restriction, the number of participants responding to the epitope that expresses the beta chain allele is indicated in parentheses; alleles present in 2 or more participants who responded to the epitope are highlighted in bold.

**Table 1:**
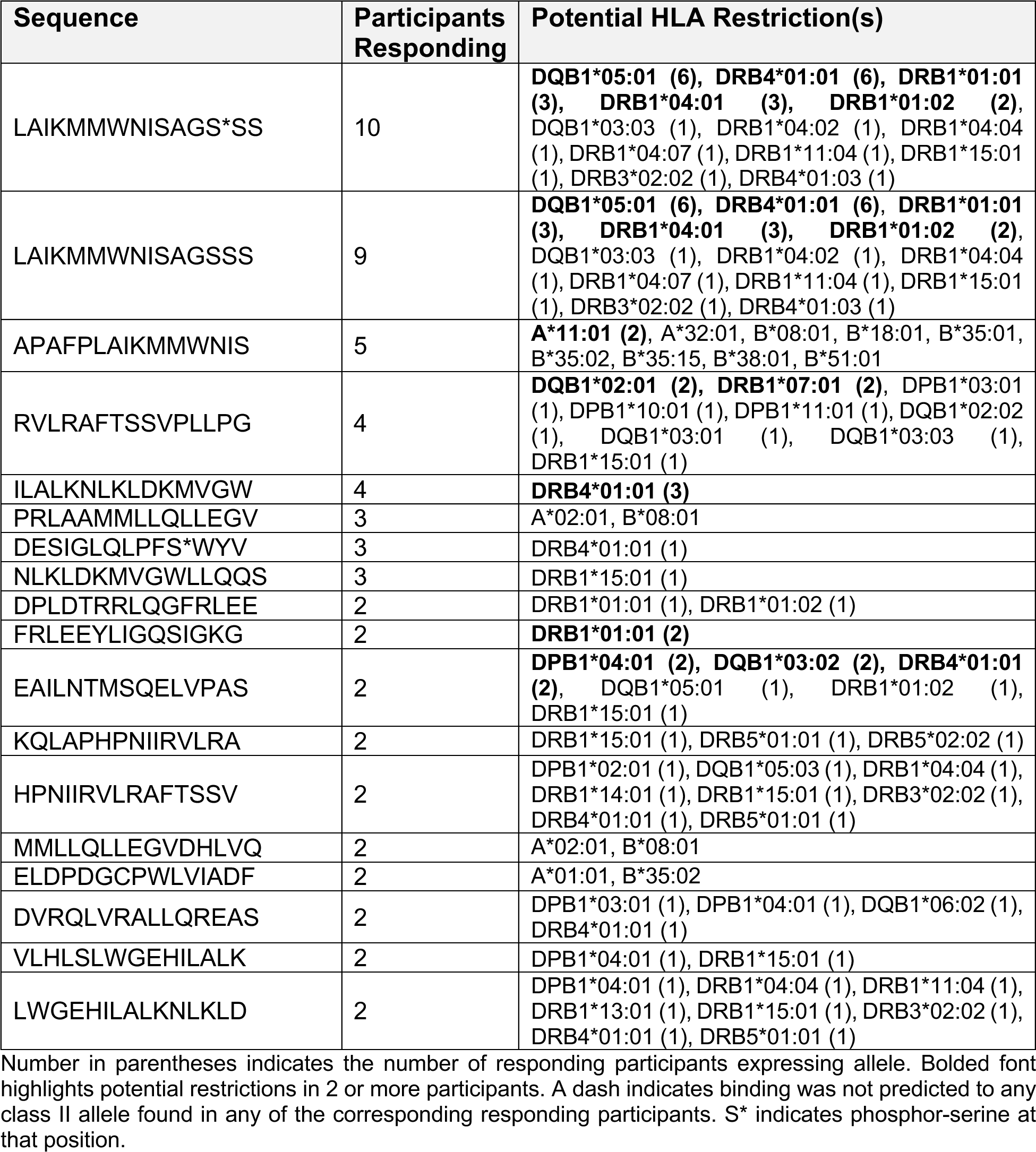
Predicted HLA restrictions for PINK1 epitopes identified in PD patients.

At least one restriction element was predicted for 14 of the 18 epitopes recognized in multiple participants. The epitopes spanning residues 216-230, in their WT and phosphorylated forms, were recognized in 9 and 10 participants, respectively, and were associated with 13 different HLA class II alleles. The most restrictions were associated with DRB1*15:01 and DRB4*01:01, which are predicted to restrict 7 and 6 of the epitopes, respectively. For the remaining four epitopes, no HLA class II restrictions were inferred. Still, in those cases, the 15-mers were found to contain 9mer/10mer peptides predicted to bind to HLA class I molecules expressed in the responding participant (*italicized* in **Table 1**), consistent with CD8 T cells corresponding to a minor fraction of the T cell expanded by the *in vitro* culture. In conclusion, multiple epitopes show promiscuous HLA binding and have several possible HLA-restrictions.

## Discussion

Identifying the specific antigenic targets recognized by T cells is essential to understand the pathogenesis of infectious disease (35, 36) and autoimmunity (37, 38). In disorders with autoimmune features, antigen identification is important for determining the basis of vulnerable cell populations, sources of antigenic substrates, and potential immune biomarkers related to the disease. In PD, a key topic of interest has been identifying the roles that T cells play in the development and/or progression of PD neurodegeneration. Our previous work (16, 17), and the work of others (18, 39) have shown that α-syn is a target of peripheral T cell responses in some PD patients. However, not all PD patients possess these auto-inflammatory T cells, and for those who do, their frequency wanes over the course of the disease (16). We have also previously shown that tau is recognized by T cells broadly in the population irrespective of age and disease status (40).

Proteins related to neurodegenerative diseases have long been examined for their possible roles in pathogenesis, and particularly recently, their roles have been expanded into non-neuronal populations such as glia, microglia (10, 41, 42), and peripheral immune cells (10, 15, 43, 44). Studies in mouse models of PD have implicated mitochondrial proteins as potential antigens, particularly the mitochondrial matrix protein, OGDH, which is implicated in the autoimmunity-linked disorder primary biliary cholangitis (11, 45). This, in turn, could be linked to the PINK1-parkin interactions that have been considered to control the turnover of damaged mitochondria in macroautophagy (31, 32), or via the formation of mitochondria-derived vesicles that can elicit a process that has been called “mitochondrial antigen presentation” (46).

Here, we tested whether OGDH and other proteins involved in the PD disease process (i.e., PINK1, PARKIN, GBA, SOD1, LRRK2) may elicit T cell responses from individuals with PD and identified the mitochondria-associated protein PINK1 as an autoantigen recognized by T cells from PD patients. The PINK1-specific T cell responses were predominantly detected in male PD patients compared to female patients. The increased incidence of PD among males is well known (47), while more recently, distinct differences in the clinical phenotype, progression, and therapeutic treatment between biological sexes have been appreciated (48). Such sex-based distinctions appear to extend to the immune system of PD, with the monocyte profile in females with PD being more inflammatory than males with PD (49). Additionally, levels of plasma cytokines have been found to differ between males and females, with significantly increased IL-4 and IL-10 levels in males with PD (50). In this study, we expand on this sex-specific immune profile with our observation of male-driven PINK1 and parkin T cell responses, as well as an IFN6FE;-bias among females in T cell reactivity towards the tested neuroantigens. These differences may be driven by the known hormonal, genetic, and environmental factors previously shown to be influenced by biological sex in the pathobiology of PD.

We identified 34 individual PINK1 epitopes responsible for mediating most of the PINK1-specific T cell response. PINK1 contains 4 major domains: the mitochondrial targeting region (aa 1-76), a transmembrane segment (aa 95-111), a protein kinase domain (aa 156-511), and a conserved C-terminal region (aa 517-581). Interestingly, we observed “regions” of reactivity in PINK1 (similar to what can be observed in pathogen (26, 36) and autoimmune (51, 52) antigens, including α-syn (53)), with distinct clusters of antigenicity in both the protein kinase domain, as well as the conserved C-terminal region. The protein kinase domain contained the most commonly reactive PINK1 epitope aa216 LAIKMMWNISAGSSS and its phosphorylated version (phosphorylated serine at aa228), which were generally both recognized by T cells from the same participant. Ser-228 is a key regulatory phosphorylation site in the kinase catalytic activity of PINK1 (54). Interestingly, the other phospho-antigen we tested, aa391 DESIGLQLPFSXWYV (Ser-402) located in the C-terminal domain, was also reactive but not its un-phosphorylated counterpart. Ser-402, like Ser-228, is a key regulatory site for the kinase activity of PINK1 (54). We hypothesize that PINK1 may be recognized as an autoantigen in PD because, similar to α-syn, it can be found within Lewy bodies (20, 21) and thus potentially phagocytosed and presented by either microglia or other CNS antigen-presenting cells to T cells (55).

As deficiencies in PINK1 and parkin activity lead to mitochondrial antigen presentation in mouse models (11), it is interesting to speculate on how this could occur in PD. PINK1 is thought to be constantly produced by local translation in axonal mitochondria and be tethered to mitochondria by the proteins synaptojanin 2 and synaptojanin 2 binding protein (32). If PINK1 is not continuously degraded, it may overstabilize parkin and block the normal mitochondrial turnover (56). It may be that PINK1, which is not normally turned over by the proteasome, macroautophagy, or chaperone-mediated autophagy, produces antigenic epitopes, including peptides with post-translational modifications such as phosphorylated residues. If the resulting PINK1-derived peptides can act as “neoantigens” that are not recognized as “self” they may activate T cell responses, a feature that occurs with immune responses to synucleins (55). Broadly, a role for altered protein degradation for PINK1 would be analogous to the blockade of protein degradation for other PD-related proteins, including α-syn (57) and modified α-syn (58), LRRK2 (59), and GBA (60).

We identified predicted HLA restrictions for the most recognized PINK1 epitopes by PD patients. Interestingly, most restrictions were associated with the DRB1*15:01 and DRB4*01:01 alleles. DRB1*15:01 was previously found to be more common in the PD population by our group and capable of presenting certain α-syn T cell epitopes (17), and is also associated with increased Alzheimer’s disease risk (61). DRB4*01:01, to our knowledge, has not been linked to PD before but is present in a high-risk haplotype associated with type 1 diabetes (62). While these are two of the most common HLA class II specificities in the general worldwide population, it is notable that they are not linked, with DRB1*15 alleles almost invariably associated with DRB5 alleles, and DRB4 generally associated with DRB1*04, 07, and 09. DRB1*01:02, DRB1*04:04, and DRB3*02:02 were predicted to be the next most frequently utilized alleles, associated with 4 epitopes each.

Moreover, the cytokines we observed responding to the neuroantigens tested (including PINK1) represent a potential multi-faceted immunological phenotype with both pro and antiinflammatory cell types at play. PD is a heterogeneous disease (63), and it is possible that the specific antigens recognized and cytokines produced are due to this underlying heterogeneity in the patient population.

In conclusion, our study has identified PINK1 as a common autoantigenic target of T cells in PD. These responses are predominantly associated with male PD individuals, multiple secreted cytokines towards PINK1 were observed, and specific epitopes and corresponding restricting HLA alleles are reported. These results reinforce the need for studying PD in the context of the immune system, with the goal of developing personalized immune-based therapies.

## Methods

### Study approval

All participants provided written informed consent for participation in the study. Ethical approval was obtained from the Institutional Review Boards at La Jolla Institute for Immunology (LJI; Protocol Nos: VD-124 and VD-118), Columbia University Irving Medical Center (CUMC; protocol number IRB-AAAQ9714 and AAAS1669), University of California San Diego (UCSD; protocol number 161224), and Shirley Ryan AbilityLab/Northwestern University (protocol number STU00209668-MOD0005).

### Study participants

Subjects with idiopathic PD and HCs were recruited by the Movement Disorders Clinic at the Department of Neurology at CUMC, by the clinical core at LJI, by the Parkinson and Other Movement Disorder Center at UCSD, and by the movement disorder specialists at the Parkinson’s disease and Movement Disorders program at Shirley Ryan AbilityLab. Inclusion criteria for PD patients consisted of i) clinically diagnosed PD with the presence of bradykinesia and either resting tremor or rigidity ii) PD diagnosis between ages 35-80 iii) history establishing dopaminergic medication benefit, iv) ability to provide informed consent. Exclusion criteria for PD were atypical parkinsonism or other neurological disorders, history of cancer within past 3 years, autoimmune disease, and chronic immune modulatory therapy. Age-matched HC were selected on the basis of i) age 45-85 and ii) ability to provide informed consent. Exclusion criteria for HC were the same as PD except for the addition of self-reported PD genetic risk factors (i.e., PD in first-degree blood relative). For the LJI cohort, PD was self-reported. Individuals with PD recruited at CUMC, UCSD, and Shirley Ryan AbilityLab all met the UK Parkinson’s Disease Society Brain Bank criteria for PD. Cohort characteristics is shown in **Supplemental Table 1**.

### Sex as a biological variable

Our study included both male and female participants (Supplemental Table 1). The results have been reported as an aggregate for the entire cohort, and additionally, with female and male participants analyzed separately.

### PBMC isolation

Venous blood was collected from each participant in either heparin or EDTA containing blood bags or tubes. PBMCs were isolated from whole blood by density gradient centrifugation using Ficoll-Paque plus (GE #17144003). In brief, blood was first spun at 1850 rpm for 15 mins with brakes off to remove plasma. Plasma depleted blood was then diluted with RPMI, and 35 mL of blood was carefully layered on tubes containing 15 mL Ficoll-Paque plus. These tubes were then centrifuged at 1850 rpm for 25 mins with the brakes off. The interphase cell layer resulting from this spin were collected, washed with RPMI, counted, and cryopreserved in 90% v/v FBS and 10% v/v dimethyl sulfoxide (DMSO) and stored in liquid nitrogen until tested. The detailed protocol for PBMC isolation can be found at protocols.io (https://dx.doi.org/10.17504/protocols.io.bw2ipgce).

### Antigen pools

For antigen candidates smaller than 1100 amino acids 15-mer peptides overlapping by 10 amino acids spanning the entire protein were used; PINK1 (115 peptides; UniProt ID Q9BXM7), parkin (91 peptides; O60260), OGDH (203 peptides; Q02218), GBA (106 peptides; P04062), and SOD1 (29 peptides; P00441). For LRRK2 (2527 aa in length; Q5S007), we predicted binding to HLA class II alleles using the 7-allele method (64) and selected the top 80 peptides with a median percentile score below 20. We also included peptides with a phosphorylated serine for PINK1 (aa228 and aa402), PARKIN (aa65), and SOD1 (aa99, 103, 106, and 108). For the pertussis peptide pool, we used a previously defined and characterized pool of 132 peptides (65). Peptides were synthesized commercially as crude material by TC Peptide Lab (San Diego, CA). Lyophilized peptide products were dissolved in 100% (DMSO) at a concentration of 20 mg/mL, and their quality was spot-checked by mass spectrometry. Overlapping and predicted class II peptides were combined to form antigen pools for all the respective antigens tested. Our lab routinely identifies CD4 and CD8 T cell epitopes, as well as uses existing data in the Immune Epitope Database and Analysis Resource (IEDB(66)) to develop peptide “megapools” (67). A detailed protocol for making megapools is found in the open-access publication by da Silva Antunes et al. (67) The utilization of these megapools allows the ability to test a large number of epitopes spanning multiple HLA-types. Specific sequence identities and reference numbers making up the various antigen pools used in this study can be found in **Supplemental Table 2. *In vitro* expansion of antigen-specific cells and FluoroSpot Assay**

*In vitro* expansion and subsequent FluoroSpot assay were performed as previously described in (15, 16). Briefly, PBMCs were thawed and then stimulated with neuroantigen or PT peptide pools (5 μg/mL) for 4 days. After 4 days, cells were supplemented with fresh RPMI and IL-2 (10 U/mL, ProSpec Bio), and fed again every 3 days. As described in detail at https://www.protocols.io/view/pbmc-stimulation-with-peptide-pools-and-fluorospot-bphjmj4n. After two weeks of culture, T cell responses to neuroantigen pools were measured by IFNγ, IL-5, and IL-10 FluoroSpot assay. Plates (Mabtech) were coated overnight at 4°C with an antibody mixture of mouse anti-human IFNγ (clone 1-D1K), mouse anti-human IL-5 (clone TRFK5), and mouse anti-human IL-10 (clone 9D7), all from Mabtech. 1 x 10^5^ harvested cells were plated in each well of the coated Fluorospot plots along with each respective antigen (5 μg/mL) and incubated at 37°C in 5% CO_2_ for 22 hrs. Cells were also stimulated with 10 μg/mL PHA (positive control) as well as DMSO (negative control) to assess non-specific cytokine production. All conditions were tested in triplicate. After incubation, cells were removed and membranes were washed. An antibody cocktail containing IFNγ (7-B6-1-FS-BAM), IL-5 (5A10-WASP), and IL-10 (12G8-biotin), all from Mabtech, prepared in PBS with 0.1% BSA was added and incubated for 2 hr at room temperature. Membranes were then washed again, and secondary antibodies (anti-BAM-490, anti-WASP-640, and SA-550, all from Mabtech) were then incubated for 1 hr at room temperature. Lastly, membranes were washed, incubated with fluorescence enhancer (Mabtech), and air-dried for reading. Spots were read and counted using the Mabtech IRIS system. Responses were considered positive if they met all three criteria: i) DMSO background subtracted spot forming cells per 10^6^ were ≥ 100, ii) stimulation index ≥ 2 compared to DMSO controls, iii) *p≤* 0.05 by Student’s t test or Poisson distribution test. The detailed protocol for the Fluorospot assay can be found at https://www.protocols.io/view/fluorospot-assay-bpspmndn.

### Flow cytometry

*In vitro* expanded cells were washed, counted, and plated in a 96 well plate at a density of 1 x 10^6^ cells/well. Cells were then stained with a mixture of the following antibodies: Fixable Viability Dye eFluor 506 (Thermo Fisher), CD3-AF700 (BD, RRID:AB_10597906), CD4-BV711 (BD, RRID:AB_2740432), and CD8-BV650 (Biolegend, RRID:AB_11125174) for 30 min at 4°C in the dark. Stained cells were then washed twice and resuspended in 100 µL PBS to be run on an LSR II flow cytometer (BD; a detailed protocol can be found at (https://doi.org/10.17504/protocols.io.bwu9pez6). FCS files produced from the LSR-II were then analyzed using FlowJo v10.8.2 software (Tree Star; RRID:SCR_008520; https://www.flowjo.com/solutions/flowjo).

### HLA typing

Participants were HLA-typed at the American Society for Histocompatibility and Immunogenetics (ASHI)-accredited laboratory at Murdoch University (Western Australia). Typing for class I (HLA A, B, and C) and class II (DQA1, DQB1, DRB1, DRB3, DRB4, DRB5, and DPB1) was performed using locus-specific PCR amplification of genomic DNA. Specific HLA loci were PCR amplified using sample specific MID-tagged primers that amplify polymorphic exons from class I (A, B, C Exons 2 and 3) and class II (DQA1 and DQB1; Exons 2 and 3, DRB and DPB1; Exon 2) major histocompatibility complex (MHC) genes relevant to epitope binding and presentation. Therefore, rare alleles that differ by a single nucleotide in exon 1 cannot be excluded due to missing coverage of exon 1. MID-tagged primers were optimized to minimize allele dropouts and primer bias. Amplified DNA products from unique MID-tagged products (up to 96 MIDs) were quantitated, pooled in equimolar ratios, and subjected to library preparation using NEBNext Ultra II library prep kits (New England Biolabs). Libraries were quantified using the Jetseq library quantitation kit (Meridian Bioscience) and High sensitivity D1000 screen tape on an Agilent 2200 Tapestation (Agililent) for concentration and size distribution. Normalized libraries were sequenced on the Illumina MiSeq platform using the MiSeq V3 600-cycle kit (2×300bp reads). Sequences were separated by MID tags, reads were quality-filtered, and alleles were called using an in-house accredited HLA caller software pipeline, minimizing the influence of sequencing errors. Alleles were called using the latest IMGT HLA allele database as the allele reference library. The algorithm was developed by E.J.P. and S.A.M. and relies on periodically updated versions of the freely available international immunogenetics information system (RRID:SCR_012780; http://www.imgt.org) and an ASHI-accredited HLA allele caller software pipeline, IIID HLA analysis suite (http://www.iiid.com.au/laboratory-testing/). Sample report integrity was tracked and checked using proprietary and accredited Laboratory Information and Management System (LIMS) and HLA analyze reporting software that performs comprehensive allele balance and contamination checks on the final dataset.

### Prediction of HLA restriction

Potential HLA class II restrictions were determined on the basis of MHC binding predictions. Predictions were performed using the IEDB’s analysis tools suite ((http://tools.iedb.org/main/tcell/), and the recommended (as of August 2023) NetMHCIIpan algorithm, v2023.05 EL 4.1 (https://services.healthtech.dtu.dk/services/NetMHCIIpan-4.1). Restrictions were assigned using a predicted binding percentile score threshold of ≤20%. Because HLA typing was unavailable for the DPA locus, predicted DPA/B dimer binding was based on known (typically strong) A/B haplotype linkages. Accordingly, DPA1*02:01 was assigned to DPB1*01:01, and DPA1*01:03 to all other DPB1. While both DQA and DQB loci are polymorphic, only haplotype linked (cis) A and B loci are believed to form stable dimers (68, 69). Thus, for DQB1*05 and 06 alleles, only dimers with DQA1*01 were considered when performing prediction analyses, and for DQB1*02, 03, and 04, only dimers with DQA1*02, 03, 04, 05, and 06 were considered. The HLA DRA1 locus is largely monomorphic, with the rare variants mutated outside of the surface exposed domains, and is thus not considered for binding predictions.

### Statistical Analysis

Statistical analyses were performed, and graphs were created using GraphPad Prism’s descriptive statistics, one-tailed Mann Whitney tests, one-way ANOVA with Dunnett’s multiple comparisons test, one-way Fisher exact tests, and Spearman r tests as applicable (GraphPad Prism, RRID:SCR_002798, v9).

## Acknowledgment

This work was partly supported by Aligning Science Across Parkinson’s (ASAP-000375 to CLA, DS) through the Michael J. Fox Foundation for Parkinson’s Research (MJFF). For open access, the authors have applied a CC-BY public copyright license to all Author Accepted Manuscripts arising from this submission. This work was also supported by the NIH T32AI125179 (GPW), the National Institute of Neurological Disorders and Stroke of the NIH R01NS095435 (AS, DS), and the JPB Foundation (DS). The funders had no role in study design, data collection, analysis, publication decision, or manuscript preparation.

## Author contributions

C.S.L.A., A.S., and D.S. participated in the design and direction of the study. G.P.W., T.M., J.R.L.J., J.S., and C.S.L.A. performed and analyzed the experiments. A.F., N.K.T., I.L., J.G.G., and R.N.A. recruited participants and performed clinical evaluations. S.A.M. and E.J.P. performed HLA typing. G.P.W., D.S., A.S., and C.S.L.A. wrote the manuscript. All authors read, edited, and approved the manuscript before submission.

---

**Supplemental Figure 1:**
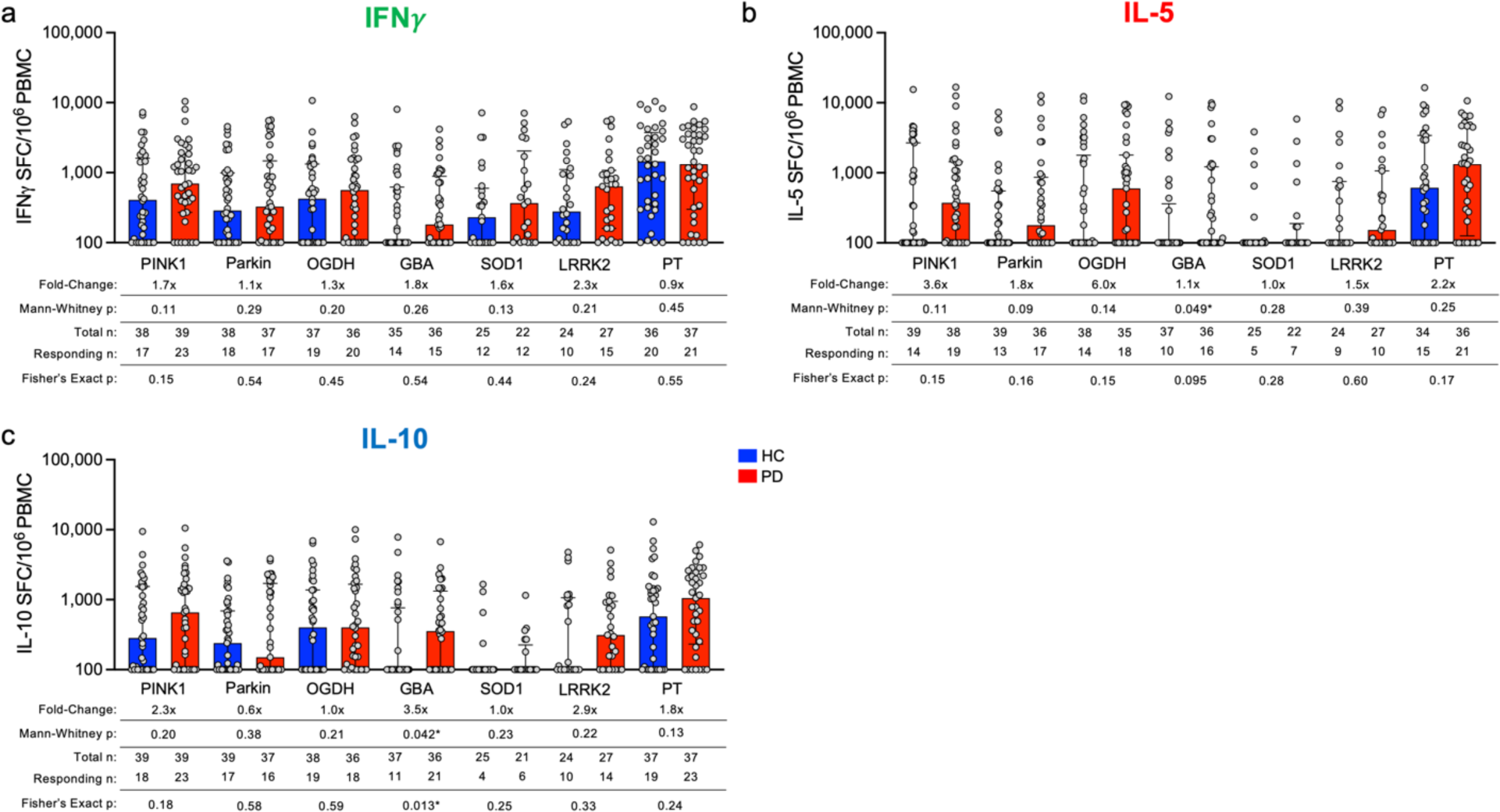
Individual cytokine responses towards neuroantigens among PD patients. The magnitude of the individual cytokine response for a) IFNγ, b) IL-5, c) IL-10 by PBMCs from PD and age-matched HC. HC (blue bars) and PD (red bars), each circle representing an individual participant. Median ± interquartile range displayed. Fold-change is in comparison to HC response. One-tailed Mann-Whitney tests were performed between HC and PD antigen-cytokine values. One-tailed Fisher tests were performed using the geometric mean of the HC group for each individual antigen as a cutoff for the test.

**Supplemental Figure 2.**
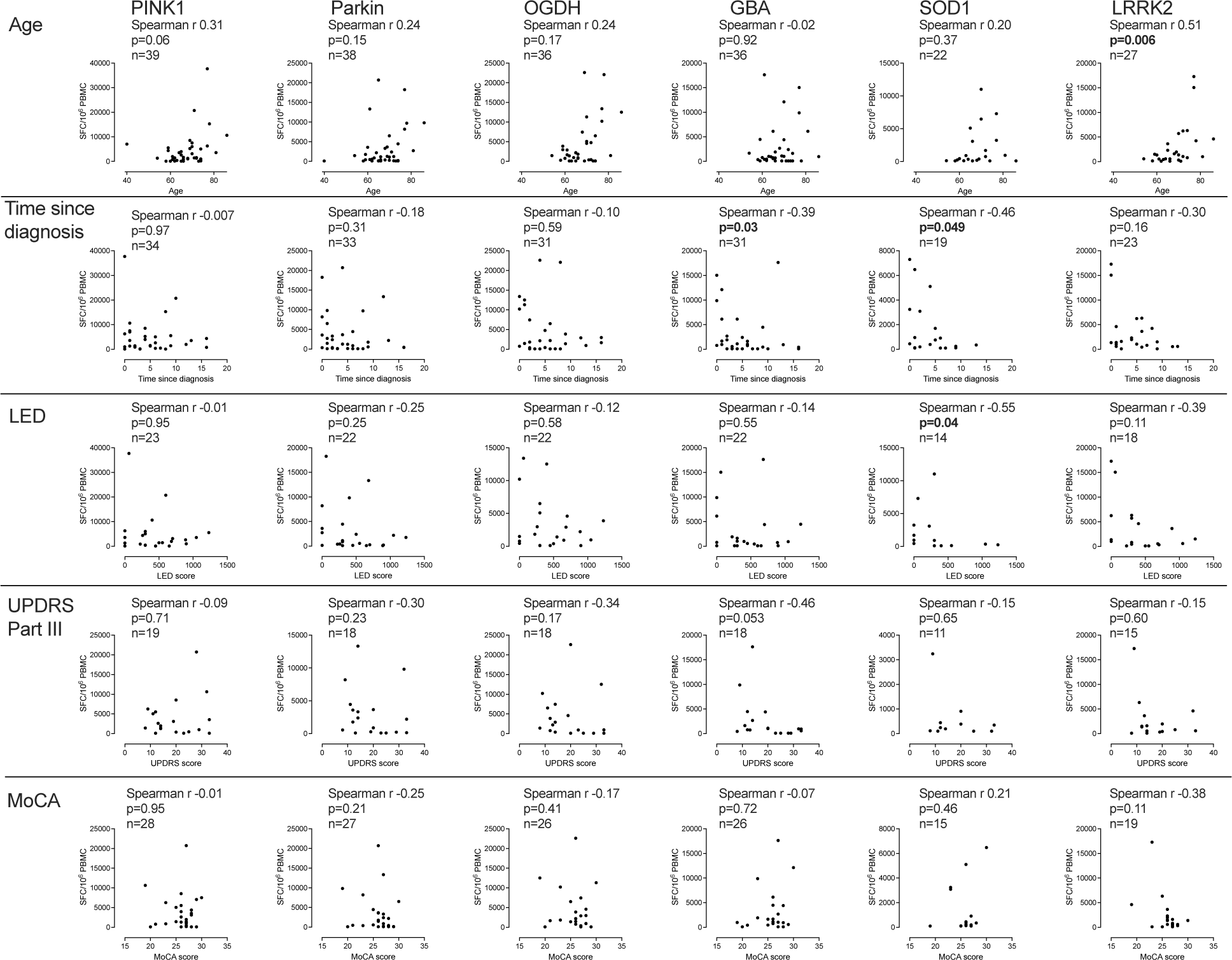
Correlation between neuroantigen-specific T cell reactivity and clinical variables. Correlation between age, time from diagnosis, LED, UPDRS part III, and T cell reactivity against MoCA and PINK1, Parkin, OGDH, GBA, SOD1, and LRRK2. T cell reactivity is the sum of the total cytokine response (IFN6FE;, IL-5, and IL-10) against the respective peptide pools as SFC per 10^6^ cultured PBMC. Correlation is indicated by Spearman r and associated p value. Each graph indicates the number of PD patients with the specific clinical variable and T cell reactivity measurement.

**Supplemental Figure 3.**
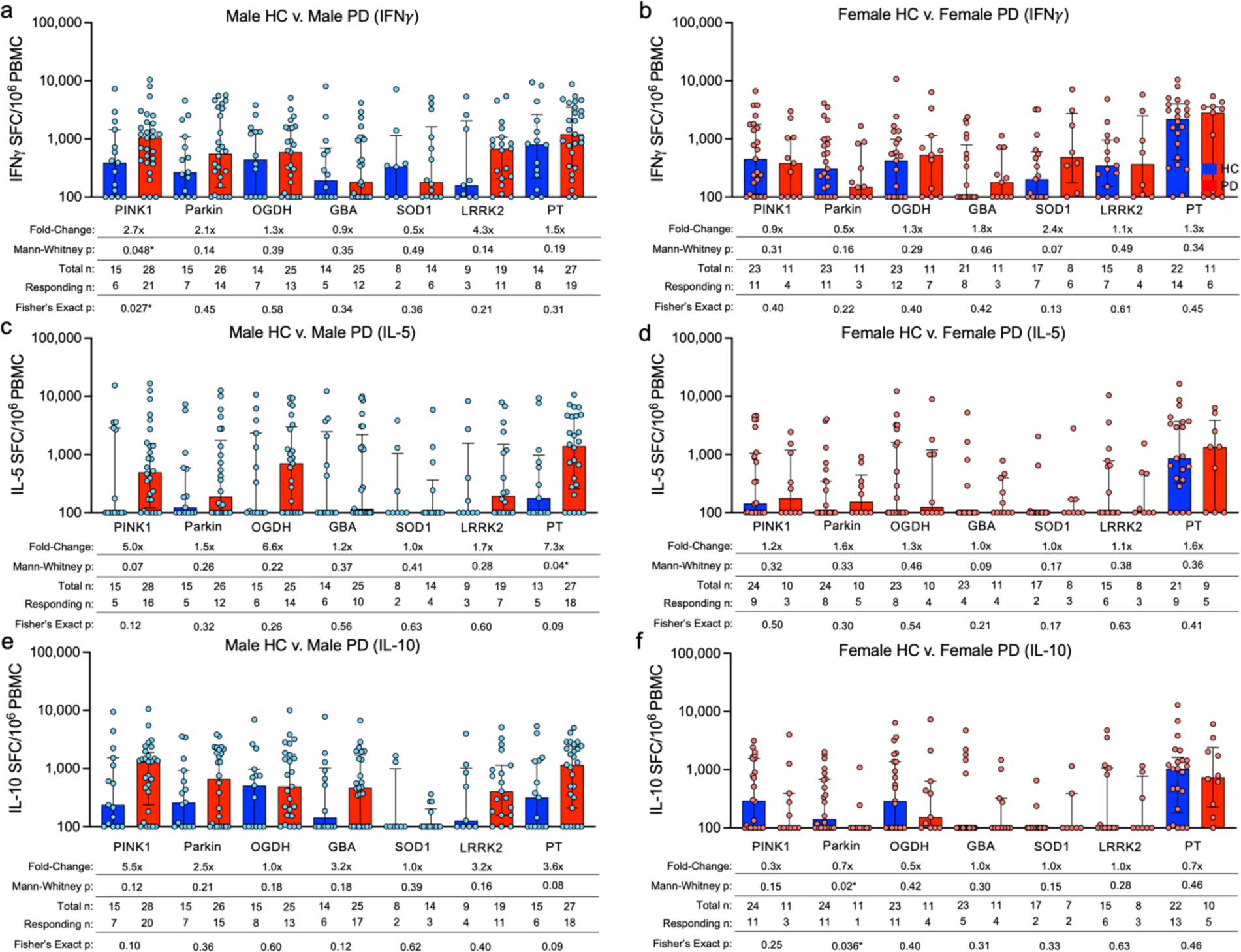
Individual cytokine responses among male and female PD patients. Magnitude of the individual cytokine response for a,b) IFNγ c,d) IL-5 e,f) IL-10 among PD and age-matched HC male and female PBMCs. HC (blue bars) and PD (red bars), each circle representing an individual participant. Median ± interquartile range displayed. Fold-change is in comparison to HC response. One-tailed Mann-Whitney tests were performed between HC and PD antigen-cytokine values. One-tailed Fisher tests were performed using the geometric mean of the HC group for each individual antigen as a cutoff for the test.

**Supplemental Table 1:**
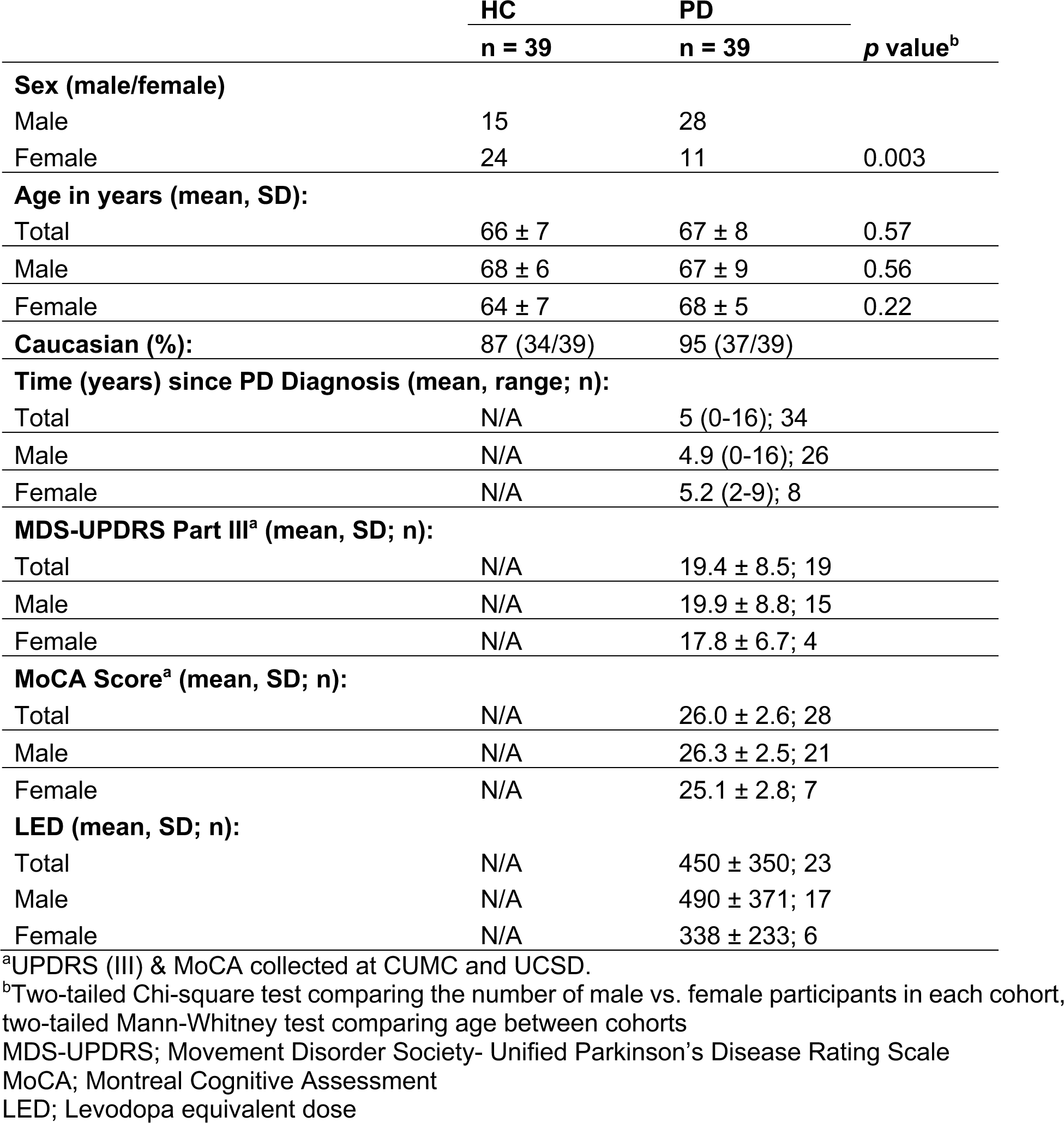
Cohort characteristics.

